# Inferring the number of spawning events from young-of-year genomic samples and otolith-derived birth dates: a richness-estimator perspective

**DOI:** 10.64898/2026.01.19.700488

**Authors:** Tetsuya Akita, Yohei Tsukahara, Hiroshige Tanaka

## Abstract

Estimating the number of spawning events per female is key to understanding individual reproductive output in batch-spawning species, yet direct observation of spawning is often infeasible in the wild. Recent advances in genetic kinship inference enable the identification of maternal half siblings from young-of-the-year genomic samples, while otolith-based age determination provides reconstruction of offspring birth dates.

Here we develop an offspring-based framework for estimating the number of clutches produced by individual females by integrating sibling structure inferred from genomic data with otolith-derived birth-date information. By recasting clutch identification as a richness estimation problem, we apply the Chao1 estimator to infer the total number of spawning events from incomplete offspring samples.

Using simulation experiments, we evaluate how sampling effort and heterogeneity in clutch size influence clutch detection and estimation. Under uniform clutch-size distributions, modest numbers of offspring sampled per maternal family (10–20 offspring) yield accurate estimates of the total number of clutches, substantially outperforming naive counts of observed birth-date classes by recovering information from rare or unobserved spawning events. In contrast, skewed or multimodal clutch-size distributions lead to underestimation at low sample sizes, indicating that uneven reproductive output increases sampling effort required for reliable inference.

Overall, our results demonstrate how offspring genomic data and otolith-derived birth dates can be jointly leveraged to reconstruct individual spawning histories under realistic sampling constraints. This perspective provides a framework for inferring within-season reproductive schedules in batch-spawning species, and highlights opportunities for integrating genomic and life-history data in fisheries monitoring and reproductive ecology.

## 1 INTRODUCTION

The number of spawning events a female completes within a single reproductive season is a fundamental aspect of fish reproductive biology but remains difficult to quantify reliably in the wild. Spawning frequency varies widely across species, from those that release a single batch of eggs to others that spawn repeatedly over extended seasons (Hunter, Lo, & Leong, 1985; Murua & Saborido-Rey, 2003). This within-season reproductive schedule shapes cumulative fecundity and patterns of energetic allocation, key components of life-history strategies (McBride et al., 2015). It can also influence population-level processes such as recruitment dynamics, which in turn affect stock assessment and management (Fitzhugh, Shertzer, Kellison, & Wyanski, 2012). Yet for many species with extended or intermittent spawning, the total number of clutches produced per season remains unknown because direct observation of spawning activity is rarely feasible under natural conditions.

Traditional approaches to estimating spawning frequency rely on repeated monitoring of captive individuals or on histological methods, such as postovulatory follicle counts or the hydrated oocyte method, that stage gonadal development (Armstrong & Witthames, 2012; Lowerre-Barbieri, Ganias, Saborido-Rey, Murua, & Hunter, 2011). Although these histological approaches have been successfully applied to large-bodied species, including Pacific Bluefin tuna (Okochi, Abe, Tanaka, Ishihara, & Shimizu, 2016; Ashida, Ishihara, Watanabe, Ohshimo, & Tanaka, 2023), their application is often constrained by limited access to reproductively important individuals, as well as by the logistical difficulty of obtaining sufficiently frequent and representative samples from highly mobile or long-lived populations (Farley, Davis, Bravington, Andamari, & Davies, 2015). Moreover, these methods often lack the temporal resolution needed to resolve discrete spawning events in batch-spawning fishes. As a result, for many ecologically and commercially important species, within-season reproductive schedules remain only partially described, limiting our ability to understand the full scope of their reproductive potential.

Advances in genetic kinship analysis and otolith microstructure now make it possible to infer reproductive processes from offspring samples alone. In many fish population surveys for stock assessment, large numbers of young-of-the-year (YOY) individuals are routinely collected because they are readily accessible. Genomic data from these samples allow the identification of sibling relationships, which form the basis for several applications in fisheries genetics, including close-kin mark–recapture (CKMR) for estimating adult abundance (Bravington, Skaug, Anderson, et al., 2016; Bravington, Grewe, & Davies, 2016; Hillary et al., 2018), inference of effective population size (Wang, 2009; Waples, 2006), and assessments of demographic connectivity (Akita, 2022; Patterson et al., 2022). Otolith microstructure from these same individuals provides daily age estimates that can be converted to individual birth dates (Campana & Moksness, 1991), and these ages also underpin high-resolution age–length keys and offer insights into growth variation under different environmental conditions (Sponaugle, 2010; García-Seoane, Meneses, & Silva, 2014; Xieu et al., 2021). When combined, these genomic and otolith data reveal whether maternal siblings were born on the same or on different days, providing a direct signal of how many times a female spawned within the season.

Here, we introduce a conceptual and statistical framework that reframes estimating the number of clutches as a richness problem. Treating each clutch as an unobserved class and each sampled offspring as an observation, we show that nonparametric richness estimators such as Chao1 (Chao, 1984) can recover the total number of clutches, including those not represented in the sample. This perspective highlights that information carried by rare birth-date classes is particularly powerful for detecting otherwise unobserved spawning events.

By leveraging existing YOY collections, this framework enables inference of within-season reproductive schedules without sampling adults or tracking individuals over time. The approach offers a simple, widely applicable tool for expanding the utility of close-kin and age-based datasets and for improving our understanding of reproductive strategies in species that spawn multiple times within a season.

Throughout this study, we use the term “number of clutches” to refer to the number of discrete spawning events completed by a female within a single reproductive season, rather than the number of eggs produced per clutch (clutch size).

## 2 ESTIMATING THE NUMBER OF CLUTCHES PRODUCED BY A FEMALE FROM THE BIRTH DATES OF HER OFFSPRING

For clarity, the primary symbols used in this study are summarized in Table 1. Figure 1 provides an overview of how maternal sibling information and otolith-derived birth dates can be combined to infer a female’s clutch structure.

**TABLE 1.** List of mathematical symbols used in the main text.

|  |  |
| --- | --- |
| $n$ | True number of clutches produced by a focal female during one breeding season. |
| $k$ | Number of clutches determined from maternal sibling relationships in the sample. |
| $s$ | Sample size of YOY individuals belonging to the focal female. |
| $f_1$ | Number of clutches observed only once in the sample (singleton). |
| $f_2$ | Number of clutches observed exactly twice in the sample (doubleton). |

**FIGURE 1.**
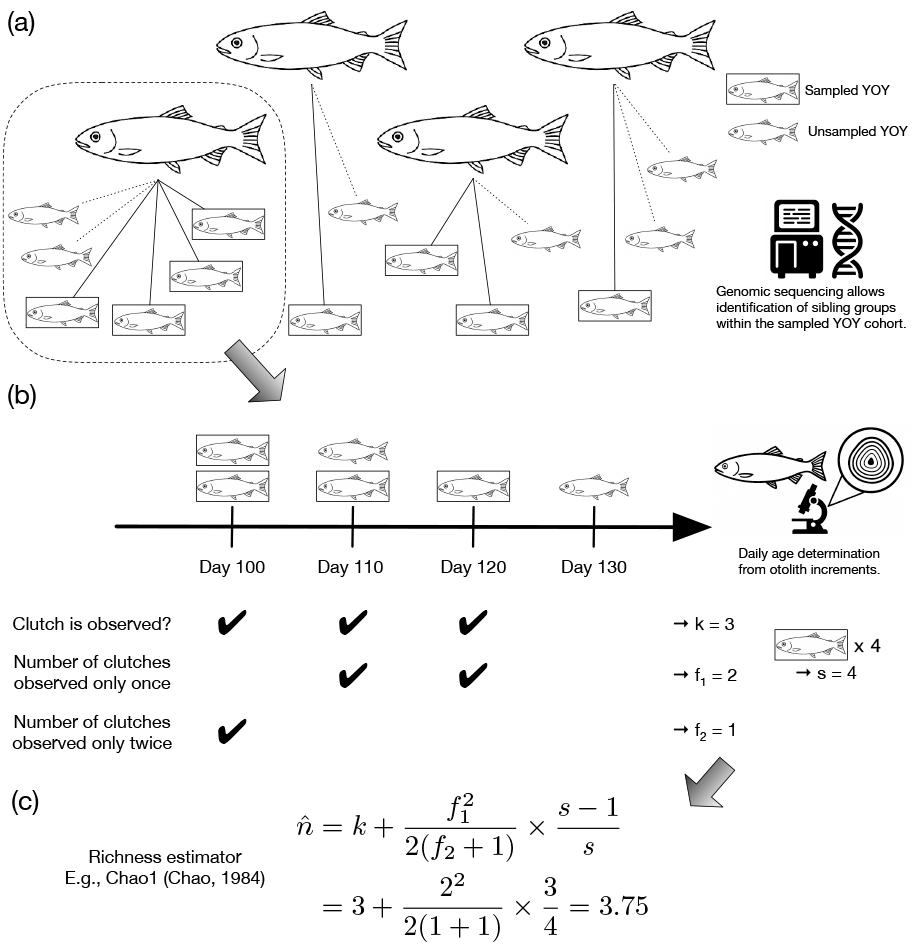
Conceptual workflow for inferring the number of spawning events from young-of-year (YOY) genomic samples and otolith-derived birth dates. (a) YOY genotypes are used to identify sibling groups without sampling parents. (b) Daily ages derived from otolith increments provide birth dates for each sibling family, revealing observed clutches as distinct birth-date clusters. (c) Applying a richness-estimator framework to each sibling family yields a lower-bound estimate of the number of spawning events, accounting for clutches not represented among the sampled YOY.

As illustrated in Figure 1, many species produce multiple batches of offspring within a single breeding season, with each spawning event forming a distinct clutch. When offspring from the same cohort can be genetically assigned to their mothers, the temporal distribution of birth dates among maternal siblings provides direct information about this within-season clutch structure. Offspring originating from the same clutch share nearly identical birth dates, whereas those from different clutches are born at distinct times.

In practice, this framework is readily applicable to young-of-the-year (YOY) samples, because the birth date of each offspring can be estimated from daily otolith increments. If two or more maternal siblings have distinct birth dates, their mother must have spawned multiple times. Consequently, the number of distinct birth-date classes observed among maternal siblings provides a lower bound on the true number of clutches produced by that female. This approach links genetic kinship and otolith information directly to the female’s reproductive schedule without requiring direct sampling or monitoring of the spawning female.

This logic naturally leads to viewing the estimation of the number of clutches as a richness problem, analogous to species richness estimation in ecology. In this analogy, each clutch corresponds to a “species” and each sampled offspring to an “individual,” and some clutches may be absent from the sample– much like rare species that go undetected in ecological surveys. This perspective allows the use of nonparametric richness estimators such as Chao1 (Chao, 1984) to infer the total number of clutches from incomplete sampling.

## 3 ESTIMATING THE NUMBER OF CLUTCHES AS A RICHNESS PROBLEM

Estimating the number of clutches can be formalized as a richness estimation problem in which each clutch represents a class that may or may not be detected among the sampled offspring of a focal female. Let *k* denote the number of distinct birth-date classes observed among the sampled siblings and let *f*_1_ and *f*_2_ denote the numbers of classes represented by one and two offspring, respectively. Because clutch sizes vary and only a subset of offspring is typically sampled, some clutches remain undetected, leading to a systematic underestimation of the true number of clutches *n* when relying solely on *k*. Let *s* denote the total number of young-of-the-year (YOY) individuals assigned to the focal female that are included in the sample.

Following the formulation in (Sard et al., 2021), which applied a finite-sample correction to the Chao1 estimator in a pedigree-based context, this structure parallels classical problems in ecological richness estimation. A widely used lower-bound estimator is Chao1, which adjusts the observed richness by leveraging the ratio of rare classes:

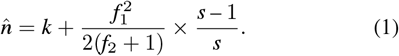

This expression requires no assumptions about the distribution of clutch sizes and provide the minimal number of clutches compatible with the observed data.

The Chao1 estimator captures a key feature of clutch detection: unequal clutch sizes create unequal detection probabilities, making some clutches far more likely to appear in samples than others. When clutch sizes are highly uneven or sample sizes are small, singletons become common and the gap between *k* and 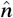 widens. Conversely, when clutch sizes are more even or sampling is intensive, *k* approaches *n* and the correction term becomes negligible. These properties make richness estimators well suited for inferring clutch structure from YOY samples when direct observation of spawning events is infeasible.

## 4 SAMPLING OFFSPRING ALONE CAN REVEAL A FEMALE’S REPRODUCTIVE SCHEDULE

To evaluate how reliably offspring-only data can recover a female’s within-season spawning schedule, we simulated YOY sampling for a female producing 15 clutches. The simulation considered three clutch-size distributions that differ in the degree and structure of heterogeneity: a uniform distribution in which all clutches contribute similar numbers of offspring, a skewed distribution in which early-season clutches are large and progressively decline in size as the season advances, and a multimodal distribution generating clusters of large and small clutches. These distributions represent biologically plausible patterns of reproductive output and determine the probability that each clutch is represented in a YOY sample (Figure 2a). To avoid unrealistically small detection probabilities, we constrained the minimum sampling probability for any clutch to be greater than 0.01.

**FIGURE 2.**
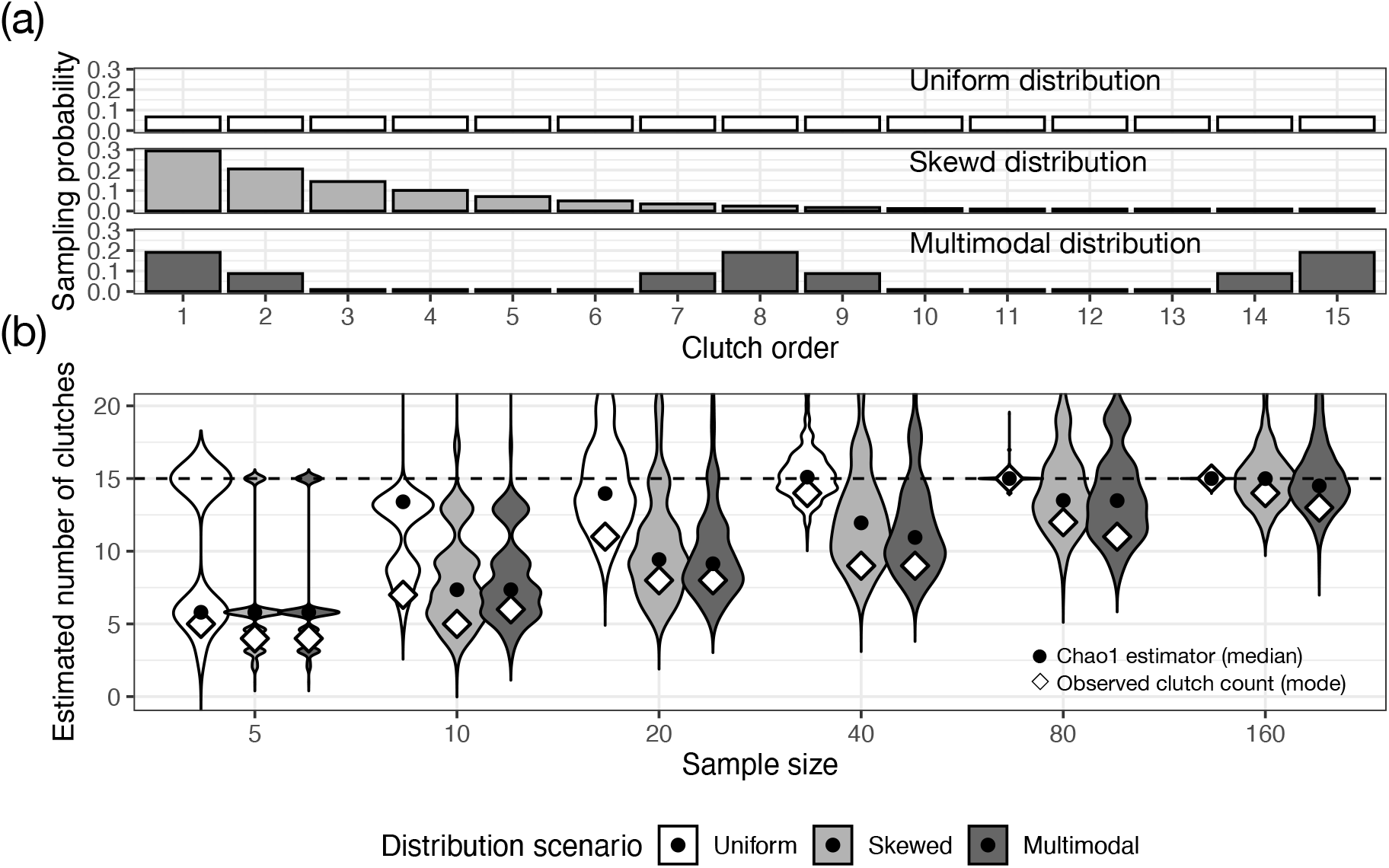
Simulation results for a female producing 15 clutches (*n* = 15), illustrating how heterogeneity in clutch size (offspring per clutch) and sampling effort influence the detection and estimation of clutches. (a) Sampling probability patterns under three clutch-size distributions–uniform, skewed, and multimodal–shown in chronological clutch order. (b) Estimated number of clutches across sample sizes from 5 to 160 for each distribution scenario. Violin plots show the distribution of Chao1 estimates based on 1000 simulation replicates, with black circles indicating median values. Diamonds indicate the most frequent observed clutch count *k* (mode across replicates). The dashed line marks the true number of clutches.

We then drew random YOY samples of increasing size (5 to 160 individuals) under a uniform clutch-size distribution and calculated both the observed number of birth-date classes *k* and the Chao1 estimator 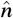.

As shown in Figure 2b, even modest sampling effort yielded accurate inference of number of clutches when all clutches contributed offspring evenly. In particular, at intermediate sampling levels (e.g., 10 to 20 offspring), 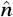 remained close to the true number of clutches and substantially outperformed *k*, demonstrating that information from rare birth-date classes can recover spawning events that are not directly observed in the sample. Similar patterns were observed when the total number of clutches was increased, with representative examples at *n* = 20 and 30 shown in Figs. S1 and S2.

When clutch sizes were heterogeneous, however, the sampling process became more challenging. Under skewed or multimodal distributions, later-season or small clutches appeared infrequently in YOY samples, reducing both *k* and 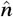 relative to the true number of clutches. Although 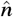 consistently exceeded *k* in these scenarios, its advantage was smaller than under the uniform distribution because a larger fraction of clutches remained entirely unsampled. These patterns emphasize that heterogeneity in detection probability–whether arising from variation in clutch size or other biological or sampling processes–can limit the recoverability of spawning events and necessitates greater sampling effort.

## 5 PRACTICAL CONSIDERATIONS FOR APPLYING THE FRAMEWORK

The clutch-based perspective developed here aligns well with many existing fish population surveys that already accumulate large numbers of young-of-the-year (YOY) individuals. Such datasets are frequently generated through YOY-only close-kin mark–recapture programs for abundance estimation as well as sibling-based analyses designed to estimate effective population size (*N*_e_ or *N*_b_). Because these approaches do not require sampling adults, the same YOY collections can be repurposed to infer within-season reproductive schedules without additional field effort.

A key practical advantage of our framework is that, once maternal siblings are available, estimating the number of clutches is relatively insensitive to moderate sampling bias, although stronger biases toward particular birth-date classes increase the sample sizes required for reliable estimation. Because each clutch is treated analogously to a species in a richness framework, moderate sampling is sufficient to recover the lower-frequency birth-date classes that correspond to rare or small clutches. In our simulations, under a uniform clutch-size distribution, even modest samples (10–20 YOY) yielded Chao1 estimates that closely tracked the true number of clutches, with representative examples at *n* = 15, 20, and 30 (Fig. 2; Figs. S1 and S2). This finding highlights the practical utility of leveraging information contained in singletons and doubletons, which contribute strongly to nonparametric richness estimators.

The method also maintains conceptual simplicity for field application. Birth dates inferred from otolith daily increments provide a direct and intuitive way to identify discrete spawning events without tracking adults or relying on continuous behavioral observations. Moreover, estimating the number of clutches does not require assumptions about the detailed distribution of clutch sizes, allowing it to be applied in situations where reproductive output is poorly characterized or highly variable among individuals or environments.

Together, these features make the framework broadly compatible with ongoing monitoring programs and demonstrate that meaningful inference on reproductive scheduling can be extracted from data sources that are already widely available but often underutilized for this purpose.

## 6 LIMITATIONS AND FUTURE DIRECTIONS FOR ROBUST APPLICATION

Although the number of clutches can be inferred from offspring data, several considerations are important for reliable application. Heterogeneity in clutch size or in detection probability can reduce the representation of late or small clutches in YOY samples, thereby lowering both the observed count *k* and, consequently, the richness-based estimate 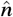 Substantially larger samples may therefore be required when reproductive output is uneven across the season or when environmental and sampling processes introduce biases in the probability that each clutch is captured.

A related practical consideration is whether a given offspring sample is likely to contain a sufficient number of maternal siblings to support inference in the first place. For an offspring sample of size *m* drawn from a single cohort, the expected number of maternal-sibling pairs is 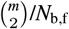, where *N*_b,f_ denotes the effective number of female breeders (Akita, 2020). This relationship highlights that maternal siblings are readily detected when the effective number of female breeders is small, either because the census size of mature females is limited or because reproductive contributions are highly skewed. Conversely, when reproductive output is more evenly distributed across many females, substantially larger offspring samples may be required to detect maternal siblings. Importantly, this consideration differs from the requirements for estimating *N*_b_, which typically rely on approximately random sampling to avoid bias in population-level inference; here, the primary objective is to obtain sufficient numbers of maternal siblings rather than to estimate a population parameter without bias. As a result, non-random or clustered sampling that increases the probability of repeatedly sampling offspring from the same mothers can be advantageous, effectively yielding larger maternal sibling groups from relatively small offspring samples.

Accurate kinship assignment is essential because the inference framework assumes that maternal sibling groups are correctly identified. Mitochondrial haplotypes provide a useful preliminary filter, as all maternal siblings must share the same haplotype. In most populations, paternal half siblings are unlikely to share mitochondrial haplotypes. However, in extreme cases where the number of spawning females is very small or mitochondrial diversity is low, paternal half siblings may share haplotypes, and genome-wide relatedness alone cannot determine whether the shared parent is the mother or the father. In addition, even with genome-wide markers, distinguishing half-sibling relationships from other close kin categories is not always straightforward, because the finite number of chromosomes limits marker independence and causes information gain to saturate as marker density increases (Tsukahara et al., 2025). As a result, some ambiguity in kinship inference is unavoidable and should be explicitly considered when applying the framework.

In the present application, this ambiguity in kinship inference is partially mitigated by the focus on young-of-the-year (YOY) individuals. Because all comparisons are made within a single cohort, avuncular relationships, which share the same expected degree of relatedness as half siblings (Trenkel, Charrier, Lorance, & Bravington, 2022), cannot arise. This cohort structure removes an important source of potential misclassification and allows more confident identification of half-sibling relationships than would be possible in mixed-age samples.

Another consideration concerns the estimation of birth dates, particularly when these are derived from otolith microstructure. Although daily increment counts are widely used, they reflect both process error, stemming from biological variability in otolith deposition (Cermeño, Uriarte, De Murguía, & Morales-Nin, 2003; Plonus et al., 2021), and observation error associated with increment interpretation (Osborne et al., 2021; Fisher & Hunter, 2018). The magnitude and structure of these uncertainties depend on species, life stage, and sampling context, and disentangling them is often difficult, especially during early life stages (Francis & Campana, 2004). Because the present framework treats birth dates as discrete temporal classes, uncertainty in age estimation can blur clutch boundaries and may lead to conservative inference of the number of clutches.

In situations where otolith-based age estimation is uncertain, unavailable, or impractical, approximate birth-date information may instead be inferred from length-based age estimates when growth trajectories are sufficiently well characterized. Such reduced-resolution approaches inevitably yield coarser temporal information and should be interpreted cautiously. Nevertheless, they may substantially broaden the applicability of the framework, particularly in large-scale monitoring programs where extensive offspring samples are available but otolith readings are limited. Quantifying the trade-offs between temporal resolution and inference accuracy, and identifying conditions under which reduced solutions remain informative, represent important directions for future work.

Finally, although this study focused on estimating the number of clutches, the framework can be extended to explore broader aspects of reproductive output. For example, variation in the relative contribution of different clutches, or temporal patterns in reproductive investment within a season, could be examined by integrating clutch-level information with additional ecological or physiological data. Estimating the number of clutches thus provides a simple and general entry point for understanding within-season reproductive dynamics in species that spawn multiple times, while highlighting opportunities for further methodological refinement and extension.

## 7 CONCLUSIONS

We introduced a clutch-based richness framework for inferring the number of spawning events from offspring-only data. By treating each clutch as an unobserved class and using birth-date variation among maternal siblings, the method enables accurate estimation of total number of clutches even when sampling is incomplete. Simulations demonstrate that modest YOY samples can recover within-season spawning patterns, particularly when reproductive output is relatively even.

Because many fish population surveys already accumulate YOY samples for purposes such as close-kin mark–recapture or estimating effective population size, the framework offers a practical opportunity to extract reproductive information from datasets that are already widely available. As such, estimating the number of clutches offers a general framework for characterizing within-season reproductive dynamics in repeatedly spawning species, using information already available from offspring samples.

## AUTHOR CONTRIBUTIONS

Tetsuya Akita and Yohei Tsukahara conceived the idea of this study. Tetsuya Akita drafted the original manuscript, developed a clutch-based richness framework, and conducted simulations. Hiroshige Tanaka provided expert input on reproductive physiology and life-history interpretation, including otolith-based age estimation and gonadal development. All authors contributed to the interpretation of the results, reviewed the manuscript draft, revised it critically, and approved the final version for submission.

## DATA AVAILABILITY STATEMENT

Code used to draw the figures can be found at https://github.com/teTUNAakita/EstNumClutches.

## SUPPORTING INFORMATION

Additional supporting information may be found in the online version of the article at the publisher’s website.

