## Supplemental Figures for "Inferring the number of spawning events from young-of-year genomic samples and otolith-derived birth dates: a richness-estimator perspective"

### Supplementary Material

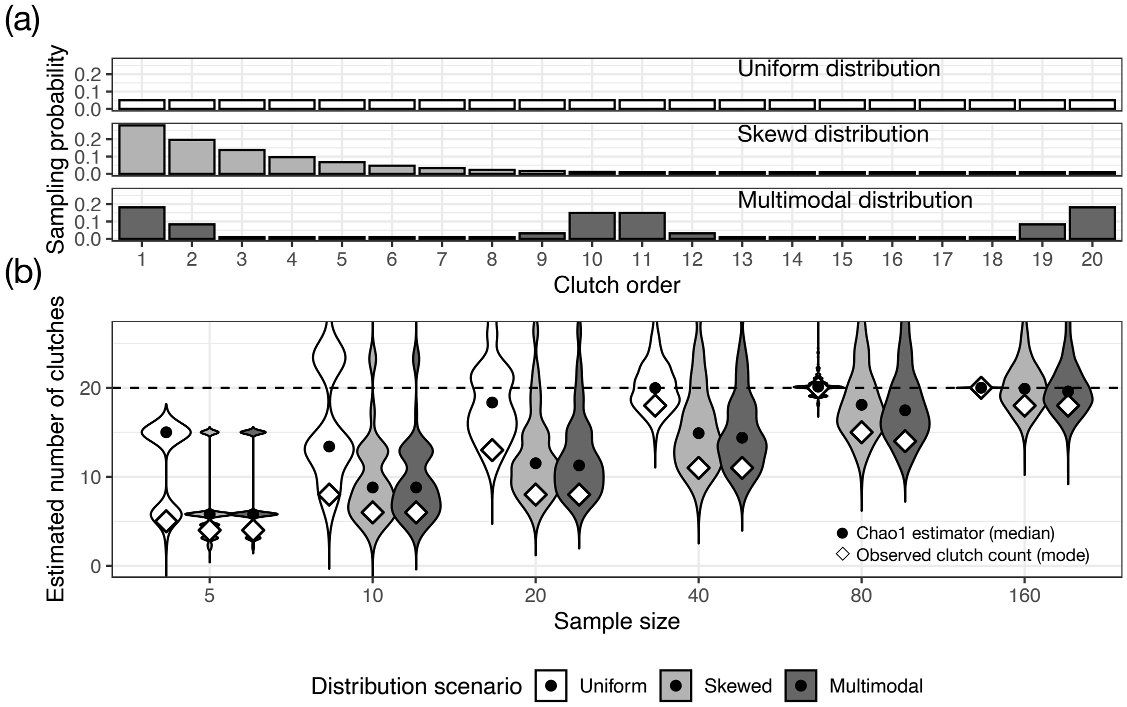


**Figure S1.** Simulation results for a female producing 20 clutches ($n = 20$), illustrating how heterogeneity in clutch size (offspring per clutch) and sampling effort influence the detection and estimation of clutches. All other conditions are the same as those in Figure 2 of the main text.

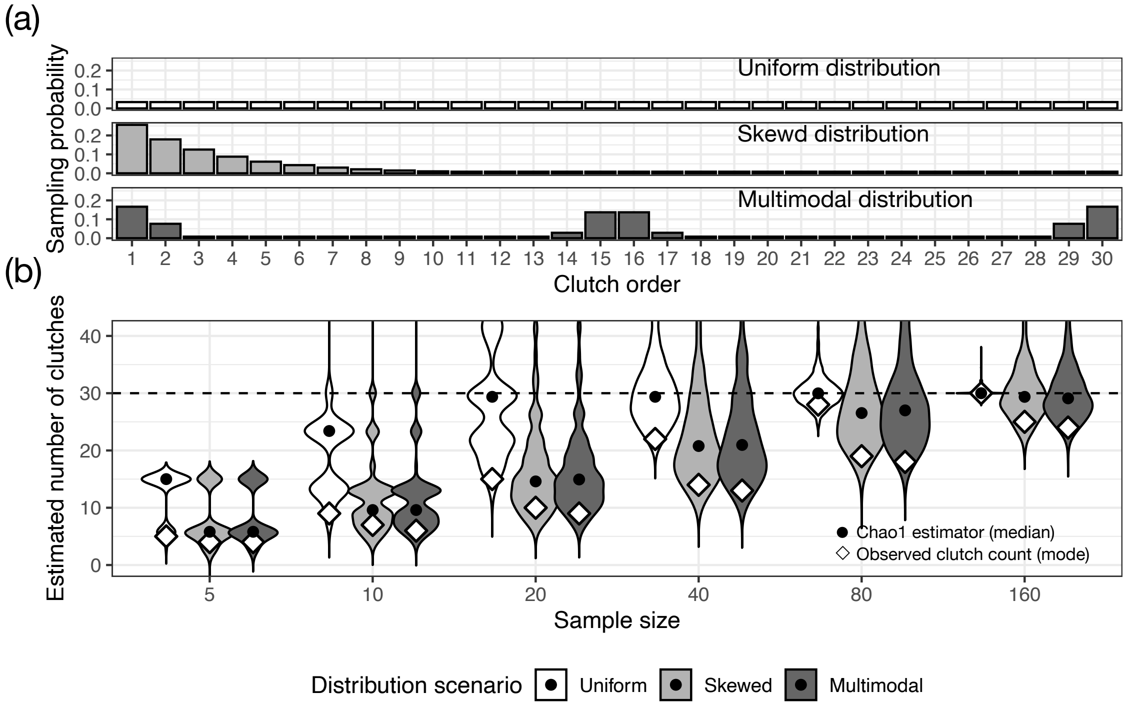


**Figure S2.** Simulation results for a female producing 30 clutches ($n = 30$), illustrating how heterogeneity in clutch size (offspring per clutch) and sampling effort influence the detection and estimation of clutches. All other conditions are the same as those in Figure 2 of the main text.
